# Genomic Response to Selection for Diurnality and Nocturnality in *Drosophila*

**DOI:** 10.1101/380733

**Authors:** Mirko Pegoraro, Laura M.M. Flavell, Pamela Menegazzi, Perrine Colombi, Pauline Dao, Charlotte Helfrich-Förster, Eran Tauber

## Abstract

Most animals restrict their activity to a specific part of the day, being diurnal, nocturnal or crepuscular. The genetic basis underlying diurnal preference is largely unknown. Under laboratory conditions, *Drosophila melanogaster* is crepuscular, showing a bi-modal activity profile. However, a survey of strains derived from wild populations indicated that high variability among individuals exists, with diurnal and nocturnal flies being observed. Using a highly diverse population, we have carried out an artificial selection experiment, selecting flies with extreme diurnal or nocturnal preference. After 10 generations, we obtained highly diurnal and nocturnal strains. We used whole-genome expression analysis to identify differentially expressed genes in diurnal, nocturnal and crepuscular (control) flies. Other than one circadian clock gene (*pdp1*), most differentially expressed genes were associated with either clock output (*pdf, to*) or input (*Rh3*, *Rh2, msn*). This finding was congruent with behavioural experiments indicating that both light masking and the circadian pacemaker are involved in driving nocturnality. The diurnal and nocturnal selection strains provide us with a unique opportunity to understand the genetic architecture of diurnal preference.

## Introduction

Although time is one of the most important dimensions that define the species ecological niche, it is often a neglected research area [1]. Most animal species exhibit locomotor activity that is restricted to a defined part of the day, and this preference constitutes the species-specific *temporal niche*. Selection for activity during a specific time of the day may be driven by various factors, including preferred temperature or light intensity, food availability and predation. The genetic basis for such phase preference is largely unknown and is the focus of this study.

The fact that within phylogenetic groups, diurnality preference is usually similar [2] alludes to an underlying genetic mechanism. The nocturnality of mammals, for example, was explained by the nocturnal bottleneck hypothesis [2], which suggested that all mammals descended from a nocturnal ancestor. Nocturnality and diurnality most likely evolved through different physiological and molecular adaptations [3]. Two plausible systems that have been targetted for genetic adaptations driven by diurnal preference are the visual system and the circadian clock, the endogenous pacemaker that drives daily rhythms. The visual system of most mammals is dominated by rods, yet is missing several cone photoreceptors that are present in other taxa where a nocturnal lifestyle is maintained [4].

Accumulating evidence suggests that diurnal preference within a species is far more diverse than previously thought. Laboratory studies [1] have often focused on a single representative wild-type strain and ignored the population and individual diversity within a species. In addition, experimental conditions in the laboratory setting (particularly light and temperature) often fail to simulate the high complexity that exists in natural conditions [1]. Furthermore, many species shift their phase preference upon changes in environmental conditions. Such “temporal niche switching” is undoubtedly associated with considerable plasticity that may lead to rapid changes in behaviour. For example, the spiny mouse (*Acomys cahirinus*) and the golden spiny mouse (*A. russatus*) are two desert sympatric species that split their habitat: the common spiny mouse is nocturnal, whereas the golden spiny mouse is diurnal. However, in experiments where the golden spiny mouse is the only species present, the mice immediately reverted to nocturnal behaviour [5].

While plasticity plays an important role in diurnal preference, there is some evidence for a strong genetic component underlying the variability seen among individuals. For instance, twin studies [6] found higher correlation of diurnal preference in monozygotic twins than in dizygotic twins, with the estimated heritability being as high as 40%. In addition, a few studies in humans have reported a significant association between polymorphisms in circadian clock genes and ‘morningness–eveningness’ chronotypes, including a polymorphism in the promoter region of the *period3* gene [7].

*Drosophila melanogaster* is considered a crepuscular species that exhibits a bimodal locomotor activity profile (in the laboratory), with peaks of activity arising just before dawn and dusk. This pattern is highly plastic and the flies promptly respond to changes in day-length or temperature simulating winter or summer. It has been shown that rises in temperature or irradiance during the day drives the flies to nocturnality, whereas low temperatures or irradiances result in a shift to more prominent diurnal behaviour [8,9]. Such plasticity was also demonstrated in studies showing that flies switch to nocturnality under moon light [10,11] or in the presence of other socially interacting flies [12].

Some evidence also alludes to the genetic component of phase preference in *Drosophila*. Sequence divergence in the *period* gene underlies the phase difference seen in locomotor and sexual rhythms between *D. melanogaster* and *D. pseudoobscur*a [13]. Flies also show natural variation in the timing of adult emergence (eclosion), with a robust response to artificial selection for the early and late eclosion phases having been shown, indicating that substantial genetic variation underlies this trait [14].

Further support for a genetic component to phase preference comes from our previous studies of allelic variation in the circadian-dedicated photoreceptor *cryptochrome* (CRY), where an association between a pervasive replacement SNP (L232H) and the phases of locomotor activity and eclosion was revealed [15]. Studies of null mutants of the *Clock* gene (*Clk*^*jrk*^) revealed that such flies became preferentially nocturnal [16], and that this phase switch is mediated by elevated CRY in a specific subset of clock neurons [17]. In other experiments, mis-expression of *Clk* resulted in light pulses evoking longer bouts of activity, suggesting that *Clk* plays a clock-independent role that modulates the effect of light on locomotion [18].

Here, using 272 natural population strains from 33 regions in Europe and Africa, we have generated a highly diverse population whose progeny exhibited a broad range of phase preferences, with both diurnal and nocturnal flies being counted. We exploited this phenotypic variability to study the genetic architecture of diurnal preference and identify loci important for this trait using artificial selection, selecting for diurnal and nocturnal flies.

## Results

### Artificial selection for diurnal preference

Flies showed a rapid and robust response to selection for phase preference. After 10 selection cycles, we obtained highly diurnal (D) and nocturnal (N) strains. The two control strains (C) showed intermediate (crepuscular) behaviour (Fig. 1). To quantify diurnal preference, we defined the ND ratio, quantitatively comparing activity during a 12 h dark period and during a 12 h light period. As early as after one cycle of selection, the ND ratios of N and D flies, and as compared to the original (control) population, were significantly different (Fig. 1A, B). After 10 generation of selection, the N and D populations were highly divergent (Fig. 1B, Table S1).

**Fig. 1.**
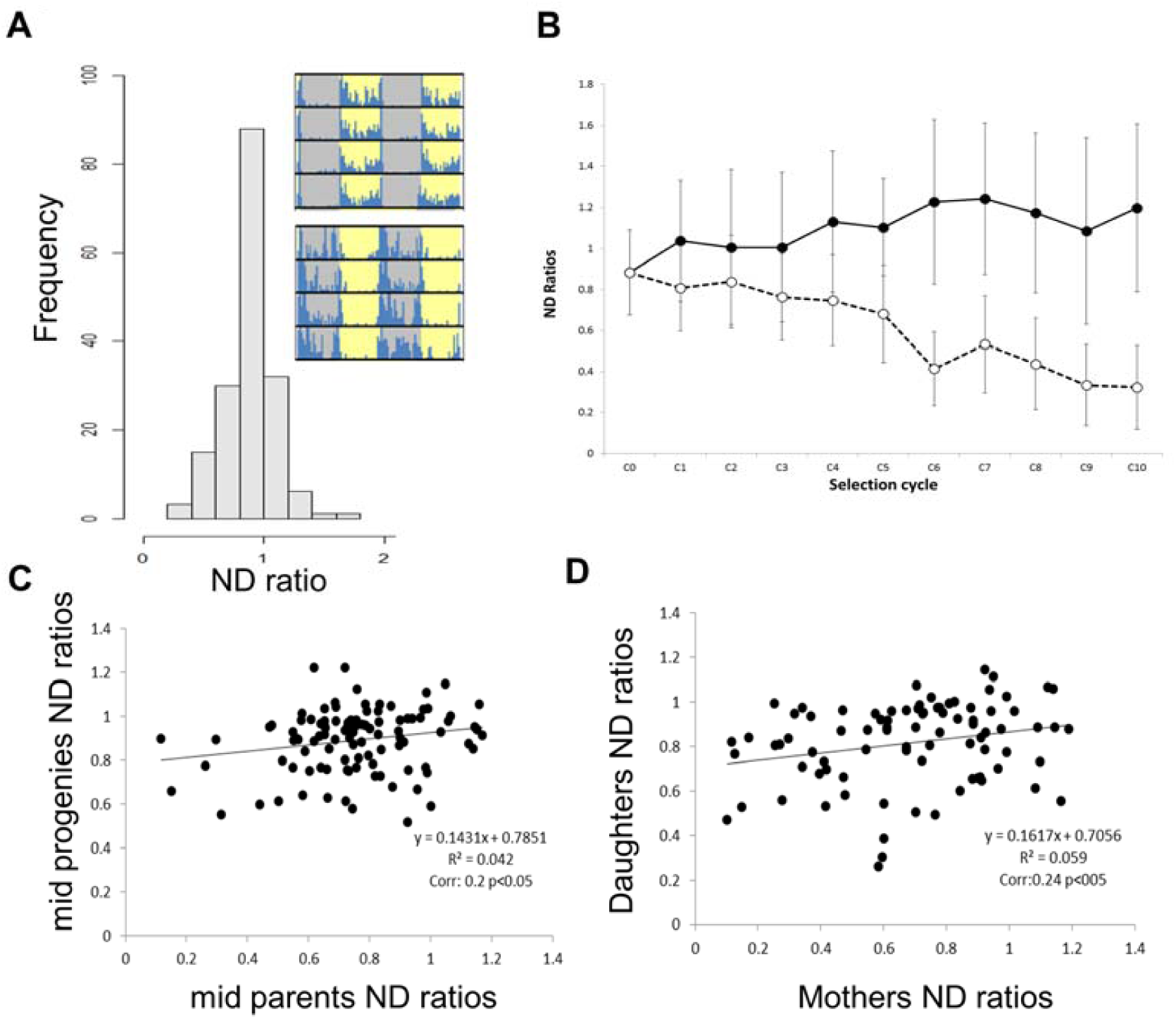
Responses to artificial selection of Nocturnal/Diurnal locomotor activity. **A.** Distribution of ND ratios of males of the starting population (C_0_, n=176). The insets show actogram examples of diurnal (top) and nocturnal flies (bottom). Grey and yellow shading represent night and day, respectively. **B.** Males average ND ratios of the selected populations per selection cycle. The black solid line represents nocturnal selection, while the dashed line represents diurnal selection. Data points correspond to average ND ratios ± standard deviation (n=104-316). The ND ratio of the original population is shown in C_0_. **C.** Correlation between mid-parents (n = 105) and mid-progenies (n =105). Correlation coefficients and p values are reported below the regression equation. **D.** Correlation between mothers (n =85) and daughters (n =85).

The estimated heritability h^2^ was higher for diurnality (37.1%) than for nocturnality, (8.4%) reflecting the asymmetric response of the two populations (Table S1). We estimated heritability by parents-offspring regression (Fig. 1C, D). The narrow-sense heritability was lower but significant (h^2=^ 14% p<0.05; Fig. 1C). The heritability value was slightly higher when ND ratios of mothers and daughters were regressed (h^2=^ 16% p<0.05; Fig. 1D) but was minute and insignificant in the case of father-son regression (Fig. S1; h^2=^ 2.5% NS).

### Effects of Nocturnal/Diurnal phenotypes on fitness

A possible mechanism driving the observed asymmetric response to selection is unequal allele frequencies, whereby a slower response to selection is associated with increased fitness [19]. We, therefore, tested whether our selection protocol asymmetrically affected the fitness of the N and D populations. After ∼15 overlapping generations from the end of the selection, we tested viability, fitness and egg-to-adult developmental time of the selection and control populations. While the survivorship of males from the three populations was similar (χ^2^= 1.6, df=2, p=0.46; Fig. 2A), we found significant differences in females. N females lived significantly longer than D females, while C females showed intermediate values (χ^2^=7.6, df=2, p<0.05; Fig. 2A). The progeny number of N females was larger than that of D females, with C females showing intermediate values (♂F_1,18_ =5.12, p<0.05; ♀F_1,18_ =5.09, p<0.05; Fig. 2B). Developmental time (egg-to-adult), another determinant of fitness, did not differ significantly between the nocturnal/diurnal populations (♂F_2,27_ =0.43, p=0.65, NS; ♀ F_2,27_ =0.27, p=0.76, NS; Fig. 2C,D).

**Fig. 2.**
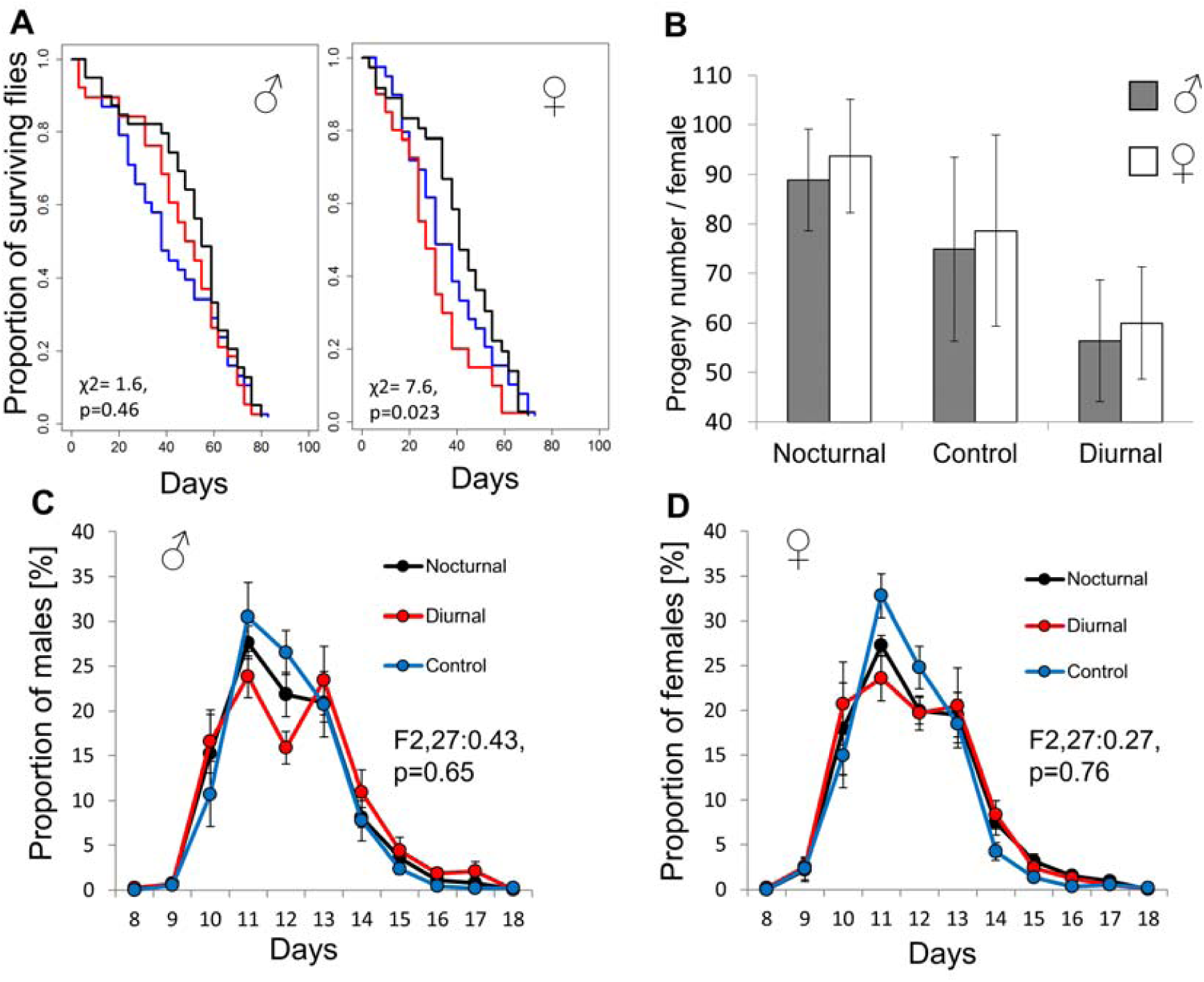
Correlated responses of fitness traits to selection. **A.** Survival curves of flies of the three selection populations (n=36-40; N: black, D: red, C: blue). The proportion of surviving flies (y axes) is plotted against the number of days (x axes). **B.** Number of progeny produced per female for N, C and D crosses. Grey and white bars indicate the number of males and females, respectively (average ± standard error). Development time (egg-to-adult) distributions are shown for male progeny (**C**) and females (**D**). Proportion (%) of progeny produced per female per day. Males: N (n=3556), D (n=2308) and C (n=2998). Females: N (n=3748), D (n=2401) and C (n=3145).

### Effects on circadian behaviour

Since the circadian system is a conceivable target for genetic adaptations that underlie diurnal preference, we tested whether the circadian clock of the N and D strains were affected by the Nocturnal/Diurnal artificial selection. Accordingly, we recorded the locomotor activity of the selection lines following three generations after completion of the selection protocol, and measured various parameters of circadian rhythmicity.

The phase of activity peak in the morning (MP) and in the evening (EP) differed between the populations (Fig. S2A). As expected, the MP of the N population was significantly advanced, as compared to that seen in both the C and D populations. The EP of the N population was significantly delayed, as compared to that noted in the two other populations. Concomitantly, the sleep pattern was also altered (S2B), with N flies sleeping much more during the day than did the other populations. While the D flies slept significantly more than C and N flies during the night, there was no difference between N and C flies.

In contrast to the striking differences seen between the selection lines in LD conditions, such differences were reduced in DD conditions (Fig. 3, Fig. S2C). The period of the free-run of activity (FRP) was longer in the C flies than in D flies, while the difference between the D and N groups was only marginally significant (Fig. 3A). No significant difference in FRP was found between N and C flies. The phases (φ) of the three populations did not differ significantly (Fig. 3B). We also tested the response of the flies to an early night (ZT15) light stimulus and found no significant differences between the delay responses (Fig. 3C).

**Fig. 3.**
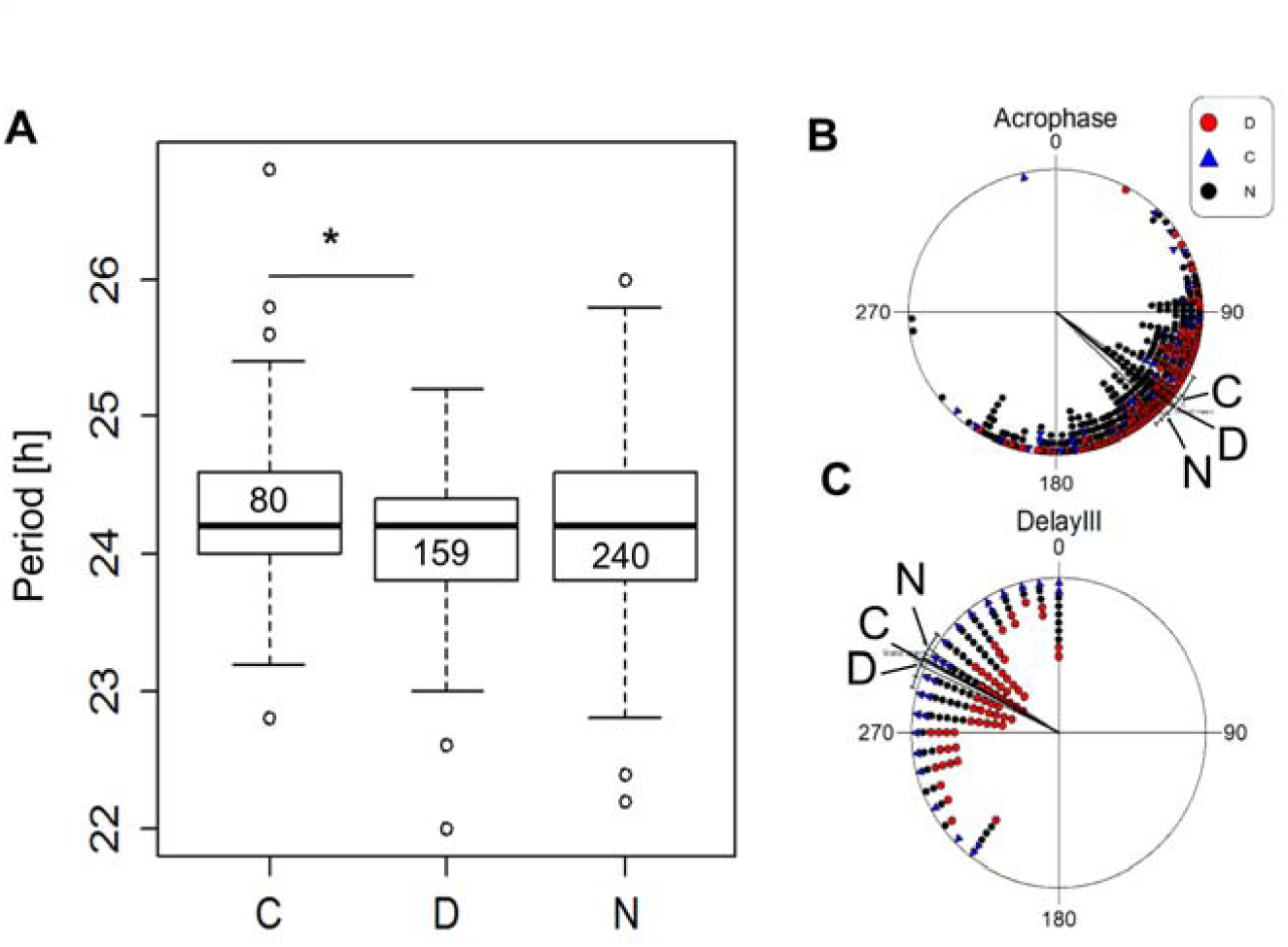
Circadian behaviour of nocturnal and diurnal selection flies. **A.** Boxplots of circadian periods under free-run conditions. Statistical differences were tested by a TukeyHSD test, with * signifying p < 0.05. **B.** Acrophase angles of the free-run activity for N (black circles, n=255), D (red circles, n=159) and C (blue triangles, n=65) populations. The free-run phases of the three populations did not differ (F_2,476_:1.91, p =0.149, NS). Lines represent mean vectors ± 95% confidence interval (CI). One hour corresponds to an angle of 15°. **C**. Phase delay angles are shown for N (black circles, n =153), D (red circles, n=155) and C (blue triangles, n=56) populations. Differences are not significant (F_2,361_ =1.47, p =0.23, NS). Lines represent mean vectors ± 95% CI.

### Circadian differences between Nocturnal/Diurnal isogenic strains

To facilitate genetic dissection of the nocturnal/diurnal preference, we generated nocturnal, diurnal and control isogenic strains (D*, N* and C*; one of each) from the selected populations. The strains were generated using a crossing scheme involving strains carrying balancer chromosomes. The ND ratios of the isogenic lines resembled those of the progenitor selection lines (Fig. S3A). The isogenic strains also differed in terms of their sleep pattern (Fig. S3B).

The circadian behaviour of the isogenic lines differed, with the N* line having a longer FRP than both the D* and C* lines (Fig. S4A). The locomotory acrophase of the N* line was delayed by about 2 h, as compared to the D* line, and by 1.38 h, as compared to the C* line (F_2,342_ =6.01, p<0.01; Fig. S4B). In contrast, circadian photosensitivity seemed to be similar among the lines, as their phase responses to a light pulse at ZT15 did not differ (F_2,359_:1.93, p =0.15, NS; Fig. S4D). Since eclosion is regulated by the circadian clock [20,21], we also compared the eclosion phase of the isogenic strains. Under LD, the eclosion phase of D* flies was delayed by ∼2 h (becoming more diurnal), as compared to both N* and C* flies, whereas no difference between N* and C* flies was detected (Fig. S4C).

### Diurnal preference is partly driven by masking

We reasoned that light masking (i.e., the clock-independent inhibitory or stimulatory effect of light on behaviour) could be instrumental in driving diurnal preference. We thus monitored fly behaviour in DD to assess the impact of light masking. We noticed that when N* flies were released in DD, their nocturnal activity was much reduced, whereas their activity during the subjective day increased (Fig. 4A). Indeed, the behaviour of N* and D* flies in DD became quite similar (Fig. 4A). Congruently, when we analysed the ND ratios of these flies in LD and DD, we found that that both N* and C* flies became significantly more “diurnal” when released into constant conditions (N*, p<0.0001; C*, p<0.001). In contrast, the ratios of D* flies did not significantly change in DD (p =0.94, NS). This result suggests that nocturnal behaviour is at least partially driven by a light-dependent repression of activity (i.e., a light masking effect).

**Fig. 4.**
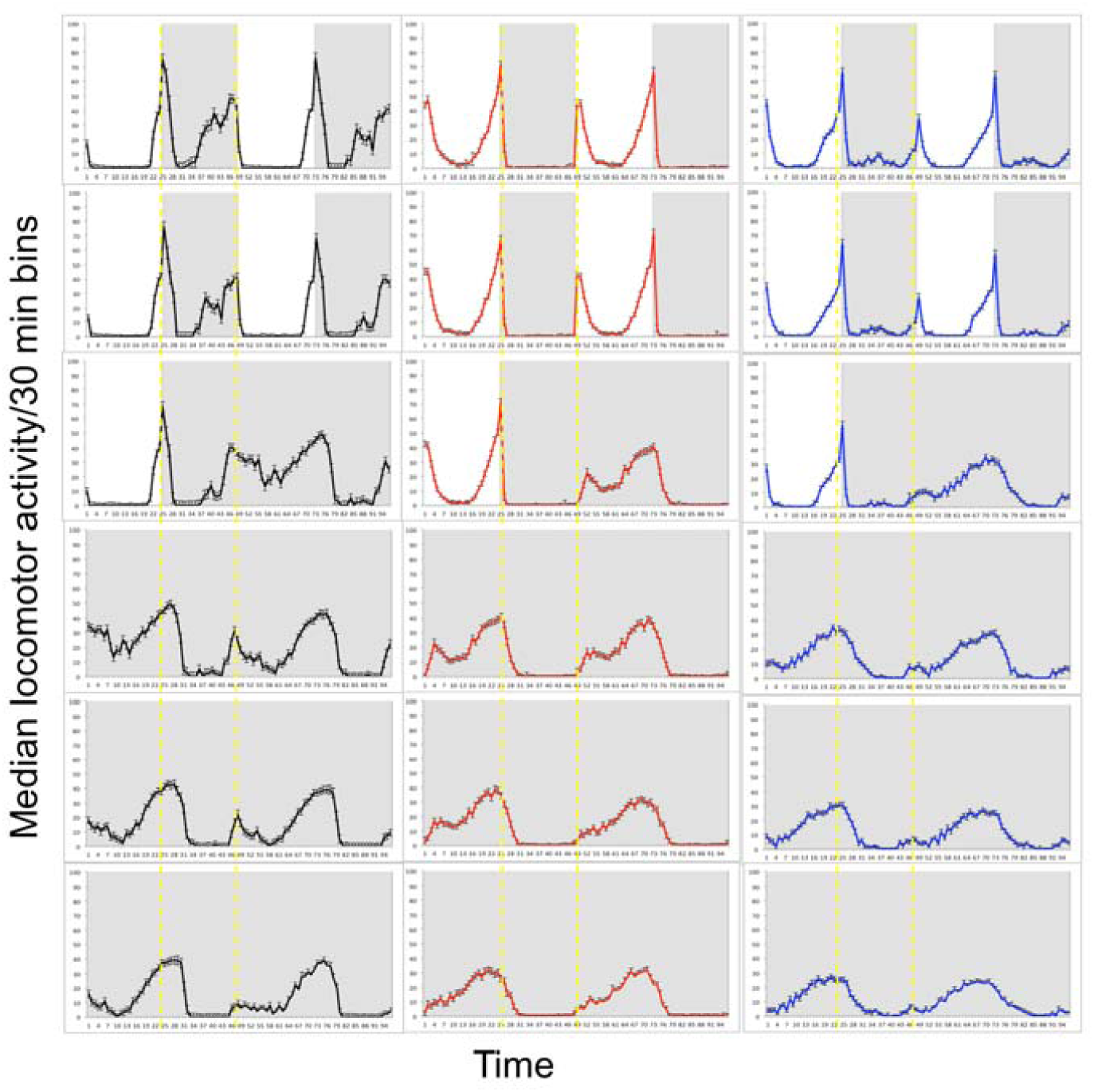
Light masking of locomotor behaviour. **A**. Double plots of the median locomotor activity (± SEM) per 30 min bin at 3 days in a 12:12 LD cycle followed by 4 days in DD. (N*, black, n=71; D*, red, n=130; C*, Blue, n =110). Shades indicate light-off. Yellow dashed lines delineate subjective nights. The overall ANOVA between 4 days in LD and 4 days in DD (starting from the second day in DD) indicates a significant effect of the light regime (i.e., LD vs. DD; F_1,594_ =105.03, p<0.0001) and genotype (F_2,594_:173.01, p<0.0001). The interaction genotype × environment was also significant (F_2,594_:102.66, p<0.0001). Post-hoc analysis (TukeyHSD text) revealed that both N* and C* flies became significantly more “diurnal” when released in constant conditions (N* p<0.0001; C* p<0.001). The ND ratios of D* flies did not significantly change in DD (p=0.94, NS).

### Correlates of the molecular clock

To investigate whether differences in diurnal preference correlated with a similar shift in the molecular clock, we measured the intensity of nuclear PERIOD (PER) in key clock neurons (Fig 5). The peak of PER signals in ventral neurons (LNv: 5^th^-sLNv, sLNv, lLNv) was delayed in N* flies, as compared to the timing of such signals in D* fly 5^th^-sLNv, sLNv and lLNv neurons. In N* and D* flies, the phases of such peaks in dorsal neurons (DN, including the clusters LNd, DN1and DN2) were similar (Fig 5).

**Fig. 5.**
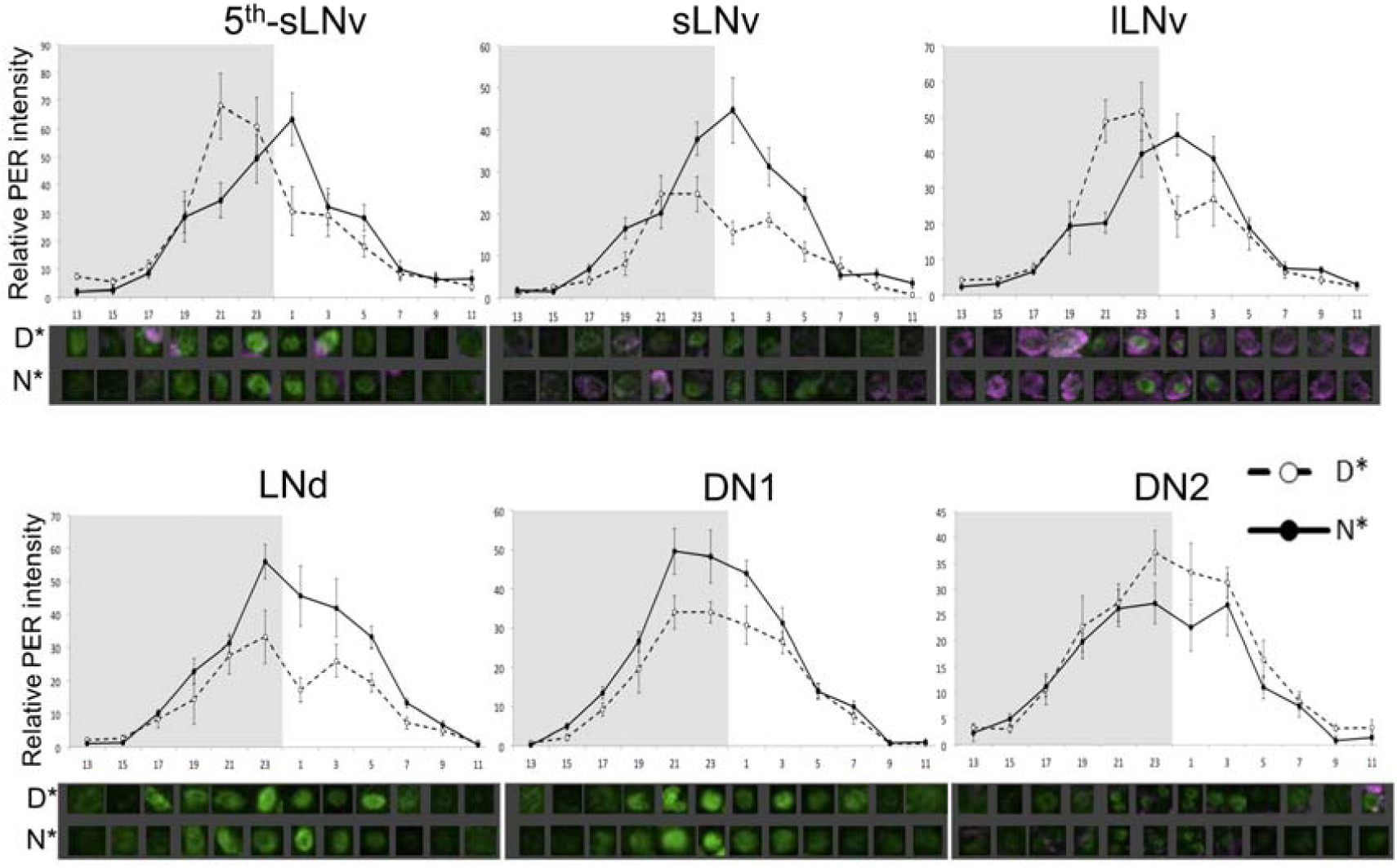
Expression of PER in clock neurons. Quantification of nuclear PER staining in N* (full lines) and D* (dashed lines) flies maintained in LD. Shaded area represents dark. Representative staining (composite, Z-stacks) is shown below each panel. Points represent averages ± standard error. The peak of PER signals in ventral neurons (LNv: 5th-sLNv, sLNv, lLNv) was delayed in N* flies, as compared to D* flies (ZT1 vs ZT21-23), as indicated by significant time x genotype interactions: 5th-sLNv χ2=12141, df=11, p<0.0001; sLNv χ2=4779.4, df=11, p<0.0001; lLNv χ2=7858, df=11, p<0.0001. The N* PER signal is stronger in sLNv (χ2=2416.6, df=1, p<0.0001), LNd (χ2=3924, df=1, p<0.0001) and DN1 (χ2=1799, df=1, p<0.0001) and weaker in DN2 (χ2=523.2, df=1, p<0.05).

We also measured the expression of the Pigment Dispersing Factor (PDF) in LNv projections (Fig. S5-S6). The signal measured in N* flies was lower than that measured in D* flies during the first part of the day (in particular at ZT3 and ZT7) yet increased during the day-night transition at ZT11 and ZT13. There were no differences seen during the rest of the night.

### Global transcriptional differences between Nocturnal/Diurnal strains

To gain insight into the genetics of diurnal preference, we profiled gene expression in fly heads of individuals from the D*, C* and N* isogenic lines by RNAseq. We tested for differentially expressed genes (DEG) in all pairwise contrasts among the three strains at two time points. We found 34 DEGs at both ZT0 and ZT12 (Table S3). An additional 19 DEG were unique to ZT0 and 87 DEG were unique to ZT12 (Table S3). Functional annotation analysis (DAVID, https://david.ncifcrf.gov/ [22]) revealed similarly enriched categories at ZT0 and ZT12 (Fig S7). The predicted products of the DEGs were largely assigned to extracellular regions and presented secretory pathway signal peptides. DEG products identified only at ZT12 were related to the immune response, amidation and kinase activity. Given the intermediate phenotype exhibited by C* flies, we reanalysed the data, searching for DEGs where C* flies showed intermediate expression (D* > C* >N* or N* > C* > D*; Table S4). The list of DEGs consisted of 22 genes at ZT0 and 62 at ZT12. Amongst the different functions represented by these new lists were photoreception, circadian rhythm, sleep, Oxidation-reduction and mating behaviour were over-represented in both D* and N* flies. For example, *Rhodopsin 3* (*Rh3*) was up-regulated in D* flies, while *Rh2* and *Photoreceptor dehydrogenase* (*Pdh*) were down-regulated in N* flies. *pastrel* (*pst*), a gene involved in learning and memory, was up-regulated in D* flies, while genes involved in the immune response were up-regulated in N* flies. The only core clock gene that showed differential expression was *Par Domain Protein 1* (*Pdp1*), which was up-regulated in D* flies. The clock output genes *takeout* (*to*) and *pdf* were up-regulated in N* flies. Overall, the results suggested that genes that are transcriptionally associated with diurnal preference are mostly found upstream (light input pathways), and downstream of genes comprising the circadian clock.

### Complementation test

We investigated the contribution of various genes to nocturnal/diurnal behaviour using a modified version of the quantitative complementation test (QCT) [23]. We tested the core circadian clock genes *per* and *Clk*, the circadian photoreceptor *cry* and the output gene *Pdf* and *Pdfr*, encoding its receptor (Fig. 6, Table S5) [24]. We also tested the ion channel-encoding *narrow abdomen* (*na*) gene, given its role in the circadian response to light and dark-light transition [25]. The QCT revealed significant allele differences in *per*, *Pdf*, *Pdfr*, *cry* and *na* (Fig. 9, Table S5), indicative of genetic variability in these genes contributing to the nocturnal/diurnal behaviour of the isogenic lines.

**Fig. 6.**
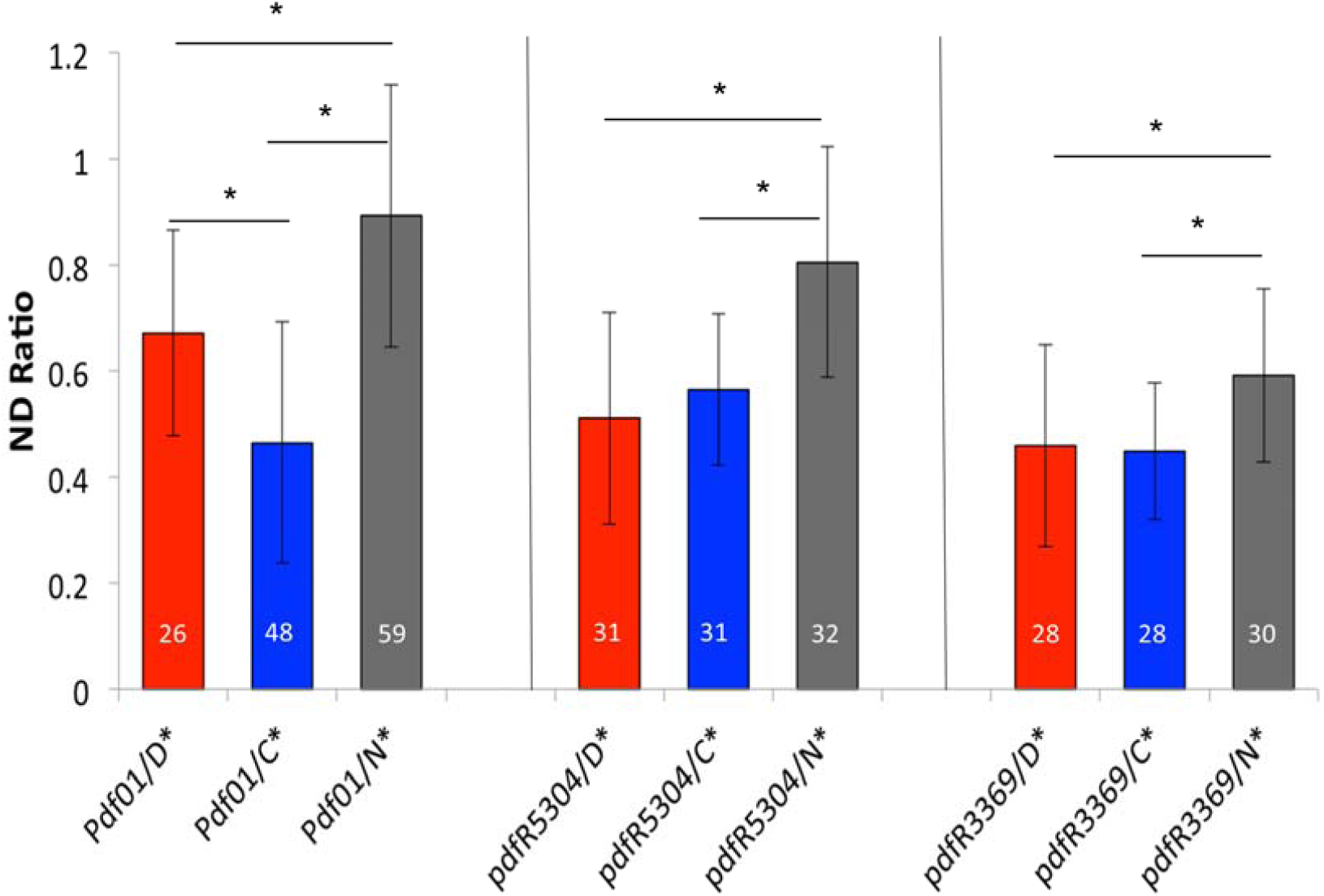
Quantitative complementation tests. Testing whether N*, D* and C* alleles vary in terms of their ability to complement the phenotype caused by the *Pdf* and *Pdfr* mutant alleles. Average ND ratio ± standard deviation of *Pdf*^*01*^ heterozygotes (left) and of *Pdfr* using p-element insertions *Pdfr*^*5304*^ (middle) and *Pdfr*^*3369*^ (right). Numbers of tested flies are reported in each chart bar. * represents p < 0.05.

Since switching from nocturnal to diurnal behaviour in mice has been shown to be associated with metabolic regulation [26], we also tested *Insulin-like peptide 6* (*Ilp6*), and *chico*, both of which are involved in the *Drosophila* insulin pathway. A significant effect was found in *Ilp6* but not in *chico* (Table S5). Other genes that failed to complement were *paralytic* (*para*), encoding a sodium channel, and *coracle* (*cora*), involved in embryonic morphogenesis [27,28].

We also tested genes that could affect the light input pathway, such as *Arrestin2* (*Arr2*) and *misshapen* (*msn*) [29]. There was a significant evidence for *msn* failing to complement but not for *Arr2* (Table S5). Various biological processes are associated with *msn*, including glucose metabolism, as suggested by a recent study [30].

## Discussion

In this study, we used artificial selection to generate two highly divergent populations that respectively show diurnal and nocturnal activity profiles. The response to selection was asymmetrical, as reflected by heritability h^2^, which was higher for diurnality (37.1%) than for nocturnality (8.4%). This may indicate that different alleles and/or different genes were affected in the two nocturnal/diurnal selections. Selections for traits affecting fitness have been shown to have higher selection responses in the direction of lower fitness [19]. This may reflect the original (natural) allele frequency, whereby deleterious traits are mostly represented by recessive alleles and favourable traits are represented by alleles at high frequencies [19]. This asymmetrical allele frequency could generate non-linear heritability, such that a slower response to selection (as seen with nocturnal flies) is associated with increased fitness [19]. Indeed, nocturnal females lived longer and produced more progeny than did diurnal females, an observation that supports a scenario of asymmetric nocturnal/diurnal allele frequencies.

To what extent is the circadian clock involved in diurnal preference? We observed that (i) PER cycling in the lateral neuron was significantly shifted in nocturnal flies, and (ii) the phase of M and E peaks in DD differed between the strains, as did their free-running period (particularly in the isogenic strains). On the other hand, our data indicate that a non-circadian direct effect of light (light masking) played a significant role in diurnal preference, particularly nocturnality, as nocturnal flies in DD conditions become rather diurnal (Fig. 5). In rodents, the differential sensitivity of nocturnal and diurnal animals to light masking has been well documented [31]. This phenomenon was observed both in the laboratory and in the field [32], with light decreasing arousal in nocturnal animals and the opposite effect occurring in diurnal animals. Light masking in flies appears to have a greater impact, as it drives flies to nocturnality.

Notably, a comparison of gene expression between diurnal and nocturnal flies highlighted just a single core clock gene *(pdp1*). This finding is reminiscent of the results of our previous study in flies [33], where transcriptional differences between the early and late chronotypes were present in genes up- and downstream of the clock but not in the clock itself. The phase conservation of core clock genes in diurnal and nocturnal animals has also been documented in mammals [34-36]. We thus suggest that selection for diurnal preference mainly targets downstream genes, thereby allowing for phase changes in specific pathways, as changes in core clock genes would have led to an overall phase change.

The main candidates responsible for diurnal preference that emerged from the current study were output genes, such as *pdf* (and its receptor *Pdfr*) and *to,* as well as genes involved in photoreception, such as *Rh3*, *TotA*, *TotC* (up-regulated in D* flies) and *Rh2* and *Pdh* (up-regulated in N*). Genes such as *misshapen (msn*) and *cry* were implicated by complementation tests. RH3 absorb UV light (λmax = 347 nm) and is the rhodopsin expressed in rhabdomer 7 (R7) flies [37], while RH2 (λmax = 420 nm) is characteristic of the ocelli [38] and *pdh* is involved in chromophore metabolism [39].

The transcriptional differences between nocturnal and diurnal flies that we identified are likely to be mediated by genetic variations in these genes or their transcriptional regulators. Our current effort is to identify these genetic variations which underlie the genetics of temporal niche preference. For this, the nocturnal and diurnal selection strains generated here will be an indispensable resource.

## Materials and Methods

The Supplemental Information has extended experimental details.

### Artificial Selection

To generate a highly genetically variable *Drosophila melanogaster* population, we pooled 5 fertilized females (4-5 days old) from 272 isofemale lines from 33 regions in Europe and Africa (Table S3) in the same culture bottle containing standard sugar food. This population was maintained at 25°C in a 12:12 LD cycle. The progeny of this population was used in the artificial selection as generation 0 (C_0_; Fig. 1). The locomotor activity of 300 males was recorded over 5 days in a 12:12 LD cycle at 25°C. Using the R library GeneCycle and a custom-made script, we identified rhythmic flies and calculated their ND ratio. In each cycle of selection, we selected 25 males with the most extreme nocturnal or diurnal ND ratios, and crossed them with their (unselected) virgin sisters.

### RNAseq

Gene expression was measured in head samples of flies collected at light-on (ZT0) and light-off (ZT12) times. Flies (3-4 days old) were entrained for 3 days in 12:12 LD cycle at 25°C at which point male heads were collected in liquid nitrogen at ZT0 and ZT12. RNA was extracted using a Maxwell 16 MDx Research Instrument (Promega), combined with the Maxwell 16 Tissue Total RNA purification kit (AS1220, Promega). RNAseq library preparation and sequencing was carried by Glasgow Polyomics using an IlluminaNextseq500 platform.

## Data Availability

The RNASeq sequencing files are available at the Gene Expression Omnibus (GEO) accession GSE116985 (use reviewer token: ibspysiwzvafhef).

